# A Potent Nonapeptide Inhibitor for the CXCL12/HMGB1 heterocomplex: A Computational and Experimental Approach

**DOI:** 10.1101/2025.02.14.638286

**Authors:** Enrico Mario Alessandro Fassi, Edisa Pirani, Valentina Cecchinato, Andrea Cavalli, Gabriella Roda, Mariagrazia Uguccioni, Jacopo Sgrignani, Giovanni Grazioso

## Abstract

Inflammation is a vital defense mechanism activated in response to injury or infection, which induces the release of cytokines and chemokines to promote tissue repair. However, persistent inflammation may result in the development of autoimmune diseases.

During pathological conditions such as rheumatoid arthritis, HMGB1 is kept in its reduced isoform and can complex with CXCL12 enhancing cell migration and exacerbating the immune responses. Small organic compounds selective for HMGB1 have been previously reported to be able to disrupt the CXCL12/HMGB1 heterocomplex, but due to their low affinity, they are unsuitable for further development as novel anti-inflammatory drugs.

We previously reported a peptide (HBP08) that binds to HMGB1 with high affinity (K_d_ = 0.8 µM), and blocks the activity of the heterocomplex, in line with a wide literature that supports the use of peptides to design protein-protein interaction inhibitors. In the present work, we computationally optimized the HBP08 peptide sequence, finding new analogues endowed with improved affinity for HMGB1. In particular, HBP08-2 inhibited the activity of the CXCL12/HMGB1 heterocomplex with an IC_50_ 10-fold lower (3.31 µM) and displayed a K_d_ 28-fold lower (28.1 ± 7.0 nM) than the parent peptide HBP08.

## Introduction

Chemokines play crucial roles in regulating the migration of leukocytes, both in normal and pathological conditions (1). These molecules signal through chemokine receptors expressed on the cell surface, which belong to the γ subfamily of rhodopsin-like G-protein-coupled receptors and represent a major target for small-molecule inhibitors, successfully employed in numerous human diseases (2). Over the past 30 years, considerable preclinical and clinical evidence has consistently supported the involvement of chemokines and their receptors in immune-related disorders (3,4). We and others have highlighted that, during inflammation, the activity of chemokines can be modulated by their interaction with other chemokines or inflammatory molecules (5–7). In particular, we have demonstrated that the alarmin High Mobility Group Box 1 (HMGB1) forms a heterocomplex with the chemokine CXCL12 (8). This interaction enhances cell migration by activating the chemokine receptor CXCR4 in the presence of a concentration of CXCL12 that would not elicit a cellular response (8).

In mammalian cells, HMGB1 is a highly conserved non-histone nuclear protein that serves as a DNA chaperon, contributing to gene transcription and DNA repair (9). In addition to its nuclear functions, HMGB1 is actively secreted under inflammatory conditions or is passively released by necrotic cells, acting as an alarmin (10,11). Structurally, it comprises two homologous but non-identical domains, BoxA and BoxB, along intrinsically disorder negatively charged C-terminal region (IDR) (12).

In the extracellular space, HMGB1 exists in various redox states, determined by the presence of an intramolecular disulfide bond between cysteines at positions 23 and 45 of BoxA. Only the reduced form of HMGB1 (fr-HMGB1), can establish a heterocomplex with CXCL12, promoting the recruitment of leukocytes to inflammatory sites via CXCR4 (8). Additionally, reduced HMGB1 can bind to the receptor for advanced glycation end products (RAGE), inducing CXCL12 secretion and autophagy (13). When oxidized by reactive oxygen species (ROS) in the extracellular space, disulfide HMGB1 (ds-HMGB1) binds to toll-like receptor 4 (TLR4) and myeloid differentiation factor 2 (MD-2), activating nuclear factor kappa-B (NF-kB) and triggering cytokine and chemokine transcription (14–16). Our previous work (17) underscored the crucial role of BoxA and BoxB in the formation of the CXCL12/HMGB1 heterocomplex. Additionally, a recent study by Mantonico et al. revealed that the IDR also plays a significant role in this protein-protein interaction (18).

HMGB1 has been identified as one of the main mediators in both acute and chronic inflammation, playing a significant role in several pathological conditions (19) including rheumatoid arthritis (20,21), systemic lupus erythematosus (22), ankylosing spondylitis (23) and other autoimmune diseases (24,25). Therefore, the identification of peptides or small molecules capable of hindering the formation of the CXCL12/HMGB1 heterocomplex could represent a novel therapeutic strategy for the treatment of the above pathological conditions. To date, only a few inhibitors of the CXCL12/HMGB1 interaction or of HMGB1 functions have been identified. Among them, glycyrrhizin, sialic acid, diflunisal, pamoic acid and a cresotic acid derivative have been shown to reduce the activity of the CXCL12/HMGB1 heterocomplex by binding to HMGB1 with an affinity ranging between 150 μM and 15 mM (8,26–30). The low affinity of the known compounds, together with their low selectivity for HMGB1 prompted us in the past to search for a compound with higher affinity for HMGB1 (31). Applying a computational pipeline, we identified a nonapeptide (HBP08) capable of binding both HMGB1 domains with a K_d_ value for the entire protein of 0.8 μM, thus representing the most potent HMGB1 binder reported to date, that significantly reduced the chemotactic activity of the CXCL12/HMGB1 heterocomplex (IC_50_ = 50 μM), without impacting HMGB1’s ability to trigger TLR4 (31). Microscale thermophoresis (MST) and nuclear magnetic resonance (NMR) experiments demonstrated that HBP08 binds both HMGB1-BoxA (K_d_ = 0.8 ± 0.3 μM) and HMGB1-BoxB (K_d_ = 17 ± 3.8 μM) with different affinity (31).

In this study, we report on the sequence optimization of HBP08, aiming at identifying new HBP08 analogues endowed with improved affinity on HMGB1-BoxB. To this aim, computational studies were accomplished to identify the most promising HBP08 analogues. Then, we fully characterized the affinity and activity of the best candidate (HBP08-2) by biophysical and biological studies.

## Results & Discussion

### Development of the HBP08/HMGB1-BoxB model

In our earlier work, we developed the HBP08 peptide to target the BoxA domain of HMGB1 (31). Biophysical assays validated our computational predictions, showing that HBP08 binds to BoxA with low micromolar affinity. However, unexpectedly, it also exhibited a K_d_ value of 17 µM for HMGB1-BoxB [29]. Moreover, combining NMR chemical shift perturbation (CSP) experiments and computer calculations, we acquired atomistic details of the interaction between HBP08 and BoxB (31).

In this study, we focus on optimizing the HBP08 sequence to design new analogues with enhanced affinity to BoxB. A long Molecular Dynamics (MD) simulation was performed on the HBP08/HMGB1-BoxB complex model (Supplementary Materials, **Figure S1**) published in our previous work (31). Cluster analysis allowed us to obtain the most representative conformation assumed by HBP08 in complex with HMGB1-BoxB during the simulation (**Figure 1A**, DOCKING-MD, purple line), which represented the starting computational model of this work (see Materials and Methods for details).

**Figure 1.**
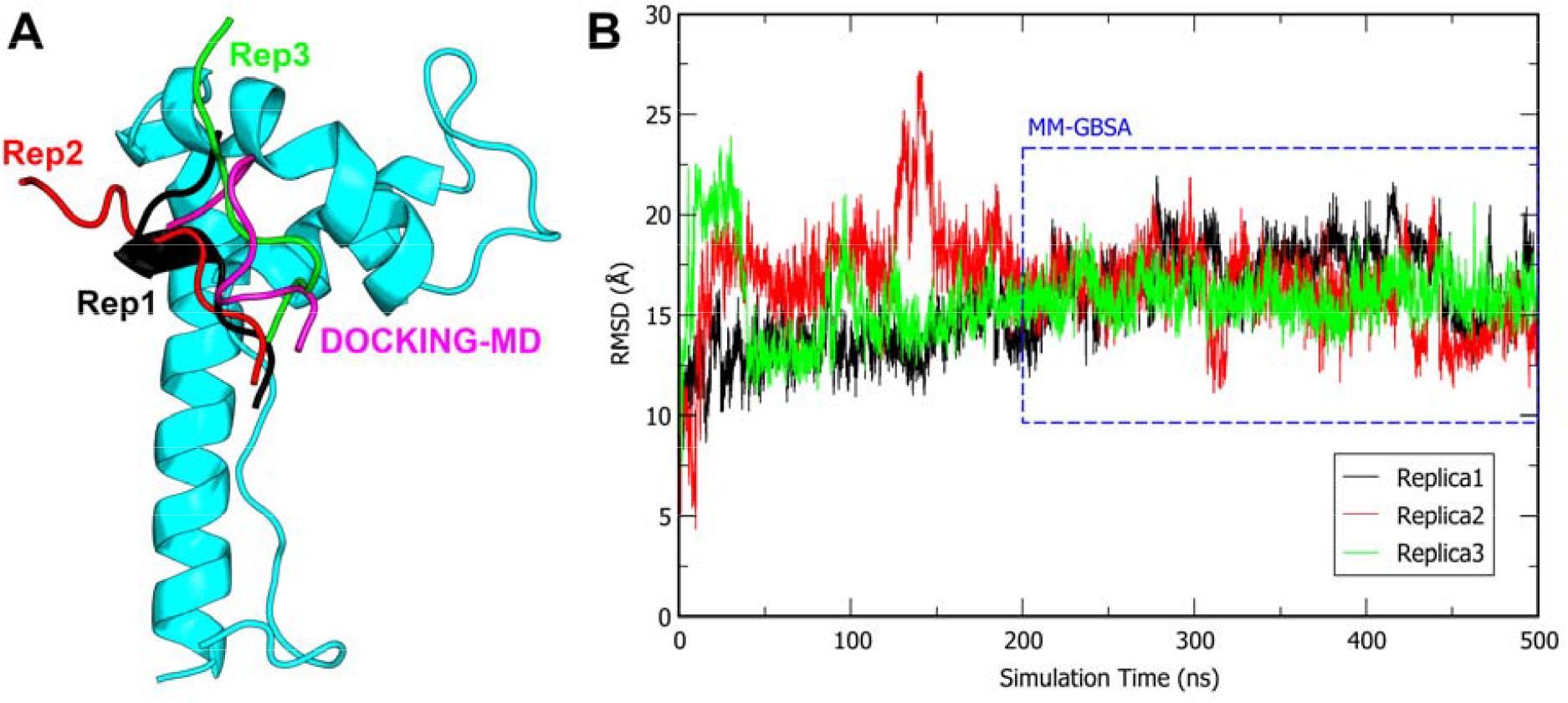
(A) Representation of the starting pose of HBP08 (magenta) and the most representative cluster conformations obtained for each MD replica (black, red and green, respectively) in complex with HMGB1-BoxB (cyan). (B) HBP08 Cα-RMSD analysis of replica1 (black), replica2 (red) and replica3 (green). The blue dotted box highlights the simulation time considered for the MM-GBSA calculations.

This complex model was then equilibrated through three independent MD simulations replicas, each lasting 500 ns, totaling 1.5 µs MD simulations (**Figure 1B**). The results of these simulations (**Figure 1**) suggested that HBP08 tends to slightly shift from the initial docking pose to more stable conformations, which were then retained until the end of the MD simulations. **Figure 1A** shows how the most representative cluster conformations of each replica shifted from the starting pose, and finally displaying a diverse HBP08 binding mode (**Figure 1A**). Consequently, Molecular Mechanics - Generalized Born Surface Area (MM-GBSA) calculations were performed on snapshots extracted from the final 300 ns of each MD replica (Supplementary Materials, **Table S1**, and **Figure 1B**) to identify the MD replica where the peptide exhibits the highest number of favorable contacts. Generally, a higher number of favorable contacts corresponds to lower ΔG values.

The third replica of HBP08 (green line, **Figure 1A**) showed the lowest ΔG value (−36.4 kcal/mol, **Table S1**) and the most stable Cα-RMSD plot (**Figure 1B**). Consequently, this was selected to perform three additional independent 500 ns-long MD simulations to better sample the conformational space of the HBP08/HMGB1-BoxB complex and to improve the robustness of the HBP08/HMGB1-BoxB model. The Cα-RMSD plots of each replica confirmed the high stability of the complex since the peptide remained stable over the whole simulation time (Supplementary Materials, **Figure S2**). Only the third replica displayed a slight variation of the binding mode after 400 ns of MD simulations, but it was noted that a binding mode close to the starting one was adopted after only 50 ns of MD simulations. MM-GBSA calculations were accomplished again on the snapshots extracted from each replica (Supplementary Materials, **Figure S2**) and the peptide ΔG values over the three replicas confirmed the reliability of HBP08-Replica3. Indeed, the average ΔG value obtained considering all the three independent replicas was −36.5 kcal/mol (Supplementary Materials, **Figure S2**), which is remarkably similar to the value attained in the first MD simulations (MD Replica3, −36.4 kcal/mol, **Table S1**). Interestingly, the binding mode of HBP08 found in the structure representative of the most populated cluster conformation of HBP08-Replica3 (Supplementary Materials, **Figure S3A**), which accounts for 89% of the explored conformational ensembles, aligns with the experimental NMR chemical shift perturbation (CSP) data from our previous work (31). In particular, HBP08 interacts with the residues A126 (hydrophobic interaction), E131 (H-bond), and N134 (hydrophobic interaction) of HMGB1-BoxB, which were involved in significant NMR chemical shifts (31). This structure constituted the starting model to design new HBP08 analogues (Supplementary Materials, **Figure S3A**).

### Computational design of HBP08 analogues

To design new HBP08 analogues, computational alanine scanning was executed on the HBP08/HMGB1-BoxB complex retrieved from HBP08-Replica3, as reported in the previous section. 500 ns-long MD simulations were performed on each Ala-mutant peptide in complex with HMGB1-BoxB. Then, MM-GBSA calculations were applied to estimate the ΔG values of each Ala-analog (**Table 1**).

**Table 1.** Binding free energy (ΔG) of the mutated peptides in complex with HMGB1-BoxB, derived from the alanine scanning calculations, subjected to 500 ns MD simulations.

| Peptide | Sequence | $\Delta G \pm SE$ (kcal/mol) |
| --- | --- | --- |
| HBP08 | GYHYERW**I**H | -36.5 |
| HBP08-Ala2 | G**A**HYERW**I**H | -35.9 ± 0.5 |
| HBP08-Ala3 | GY**A**YERW**I**H | -37.6 ± 0.4 |
| HBP08-Ala4 | GYH**A**ERW**I**H | -33.8 ± 0.4 |
| HBP08-Ala5 | GYHY**A**RW**I**H | -36.9 ± 0.3 |
| HBP08-Ala6 | GYHYE**A**W**I**H | -38.8 ± 0.4 |
| HBP08-Ala7 | GYHYER**A**I**H** | -26.5 ± 0.3 |
| HBP08-Ala8 | GYHYERW**A**H | -37.2 ± 0.5 |
| HBP08-Ala9 | GYHYERW**I****A** | -32.6 ± 0.3 |

The attained results (**Table 1**) suggested that the residue HBP08-W7 was fundamental for the interaction of HBP08 to HMGB1-BoxB, since a loss of about 10 kcal/mol in the ΔG value was observed, compared to the parent peptide. Additionally, HBP08-Y4 and HBP08-H9 were also found to be critical, as a decrease of 2.6 kcal/mol (Y4) or 3.8 kcal/mol (H9) in binding free energy for HMGB1-BoxB were observed. Therefore, residues Y4, W7, and H9 were considered *hot spots* and retained in the sequence of the new HBP08 analogues. Conversely, the other residues, poorly contributing to the binding free energy (*non-hot spots*) were systematically replaced by different amino acids by applying the “affinity maturation protocol” (32), aiming at designing new peptides with enhanced complementarity to HMGB1-BoxB. Initially, the “affinity maturation protocol” was applied on the residues at positions 2 and 6, generating a total of 20^2^ mutant peptides (i.e., 400). Upon completing the calculations, the mutant peptides were ranked according to the ΔAffinity and ΔStability values derived from the affinity maturation results (see Materials and Methods for details). Then, the best nine peptides, selected by considering the ΔAffinity, ΔStability, and both parameters (namely, mixed group) values, were simulated in complex with HMGB1-BoxB by 500 ns long MD simulations and their ΔG values were calculated by MM-GBSA approach (**Table 2**). Accordingly, the peptide containing an Arg in position 2 and a Met in position 6 (**HBP08-1**) of the sequence were considered the most promising, since it showed a predicted ΔG value about 3 kcal/mol lower than that of the parent peptide HBP08.

**Table 2.**
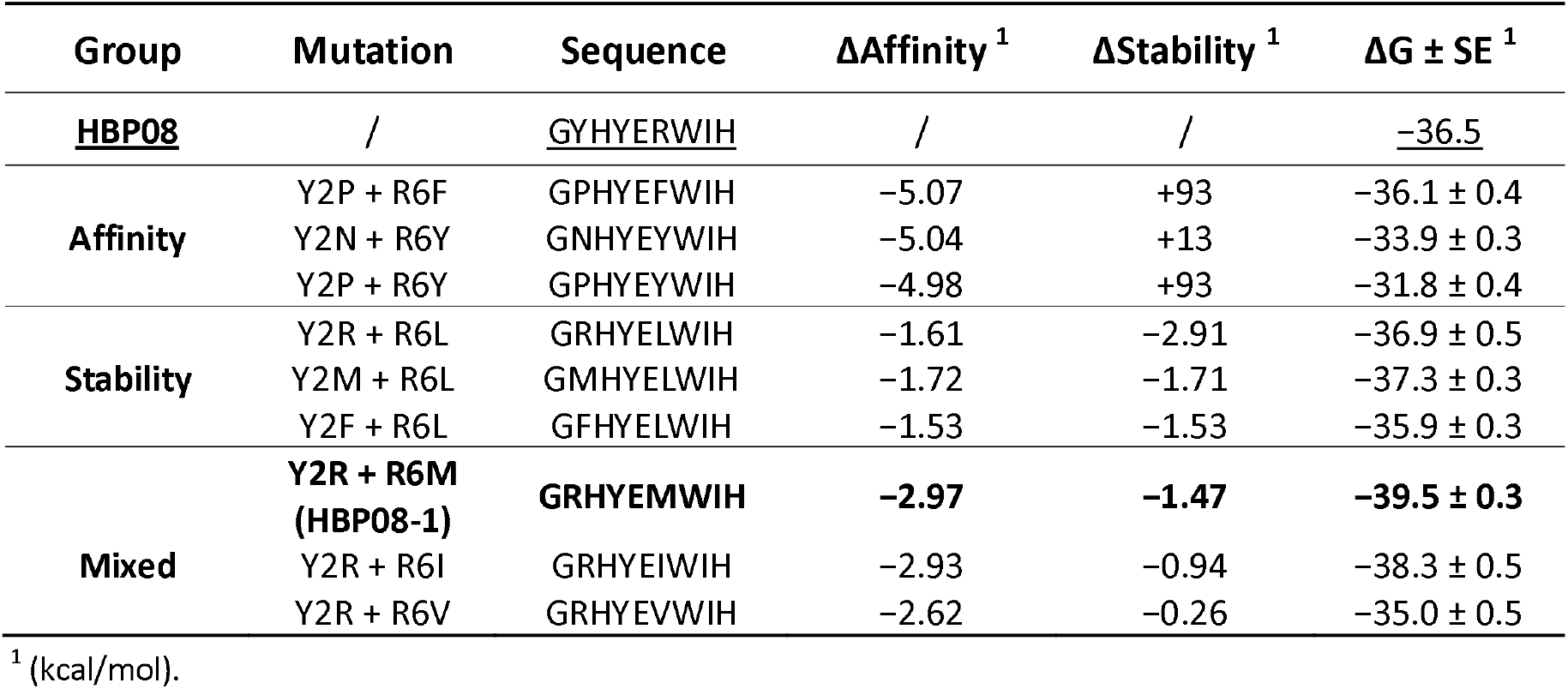
Binding free energy (ΔG) of the HBP08 mutated peptides in complex with HMGB1-BoxB, derived from the affinity maturation protocol in which Y2 and R6 were simultaneously mutated, subjected to 500 ns MD simulations.

To further corroborate these data, and to further verify the robustness of the obtained computational results, **HBP08-1** was additionally simulated in complex with HMGB1 by two additional MD replicas, attaining a predicted mean ΔG value of −38 kcal/mol (Supplementary Materials, **Table S2**), a value still lower than that of the parent peptide HBP08 (**Table 2**).

The structure representative of the most populated cluster of the **HBP08-1** peptide in complex with HMGB1-BoxB, accounting for 83% of conformational ensembles explored (Supplementary Materials, **Figure S3B**), was visually inspected and compared to the parent peptide HBP08. Interestingly, the Y2R mutation allows to create an additional H-bond with both E131 and N135 of HMGB1-BoxB compared to HBP08. Moreover, this new conformation adopted by **HBP08-1** allows its Y4 to form an extra H-bond interaction with N134, in addition to the one with E131 that can also be observed in the case of HBP08. Instead, the Ile in position 8 of **HBP08-1** is positioned in the same hydrophobic pocket surrounded by F103, A126, L129, C106, as it is observable for HBP08. Finally, the H9 side chain of **HBP08-1** slightly changes its position compared to HBP08 allowing the formation of a π-π interaction with F103 instead of R97, with which it still forms H-bond interactions (Supplementary Materials, **Figure S3B**).

This **HBP08-1**/HMGB1-BoxB model was selected for two additional steps of affinity maturation process. Specifically, H3 and I8 were simultaneously mutated to all possible natural amino acids; however, none of the new peptides showed improved predicted affinity for HMGB1-BoxB (Supplementary Materials, **Table S3**). Finally, G1 and E5 were simultaneously mutated, and the best three mutant peptides, ranked by ΔAffinity, ΔStability, and considering both parameters (namely, mixed group), were subjected to MD simulations over 500 ns, and their ΔG values were estimated by MM-GBSA approach (**Table 3**).

**Table 3.**
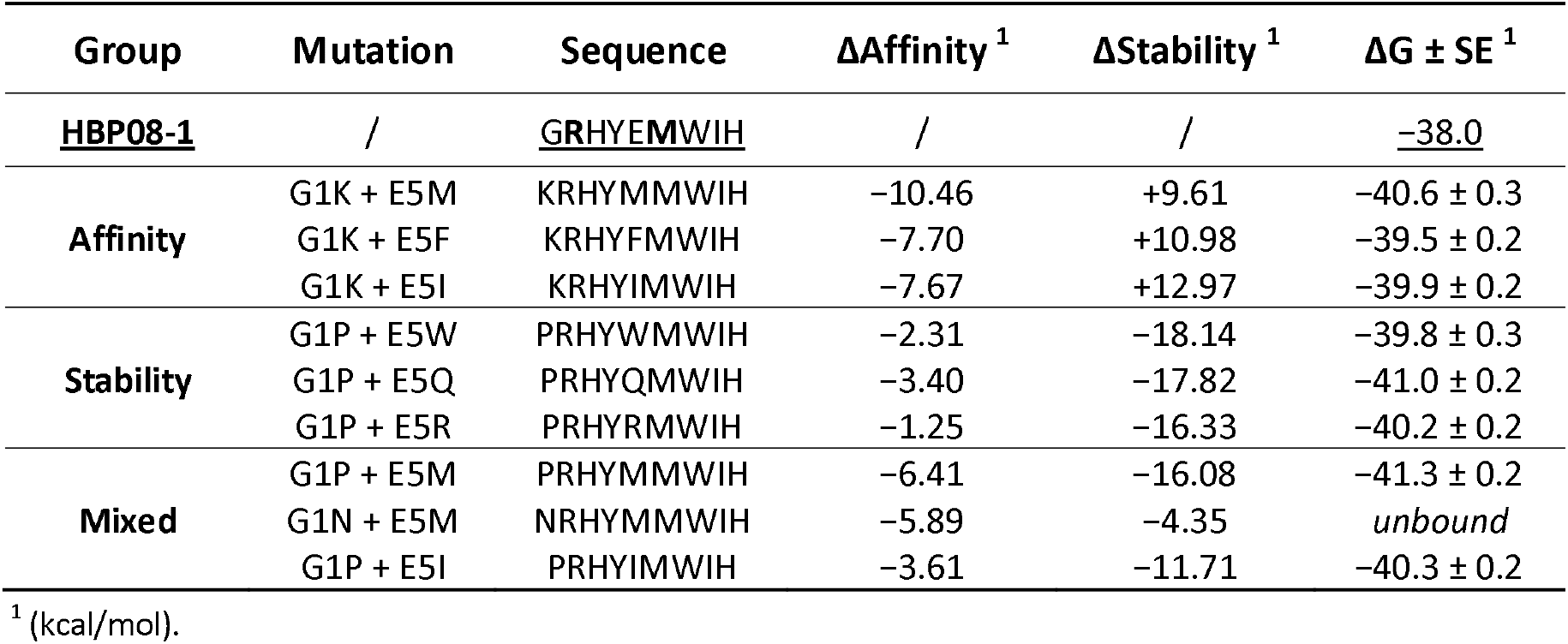
Binding free energy (ΔG) of the HBP08-1 mutated peptides in complex with HMGB1-BoxB, derived from the affinity maturation protocol in which G1 and E5 were simultaneously mutated, subjected to 500 ns MD simulations.

All mutant peptides displayed ΔG values lower that the parent peptide **HBP08-1**. The peptides showing ΔG values more than 2 kcal/mol lower were selected for two additional, and independent, 500 ns-long MD simulations replicas, to improve the robustness of the results (**Table 4**).

**Table 4.**
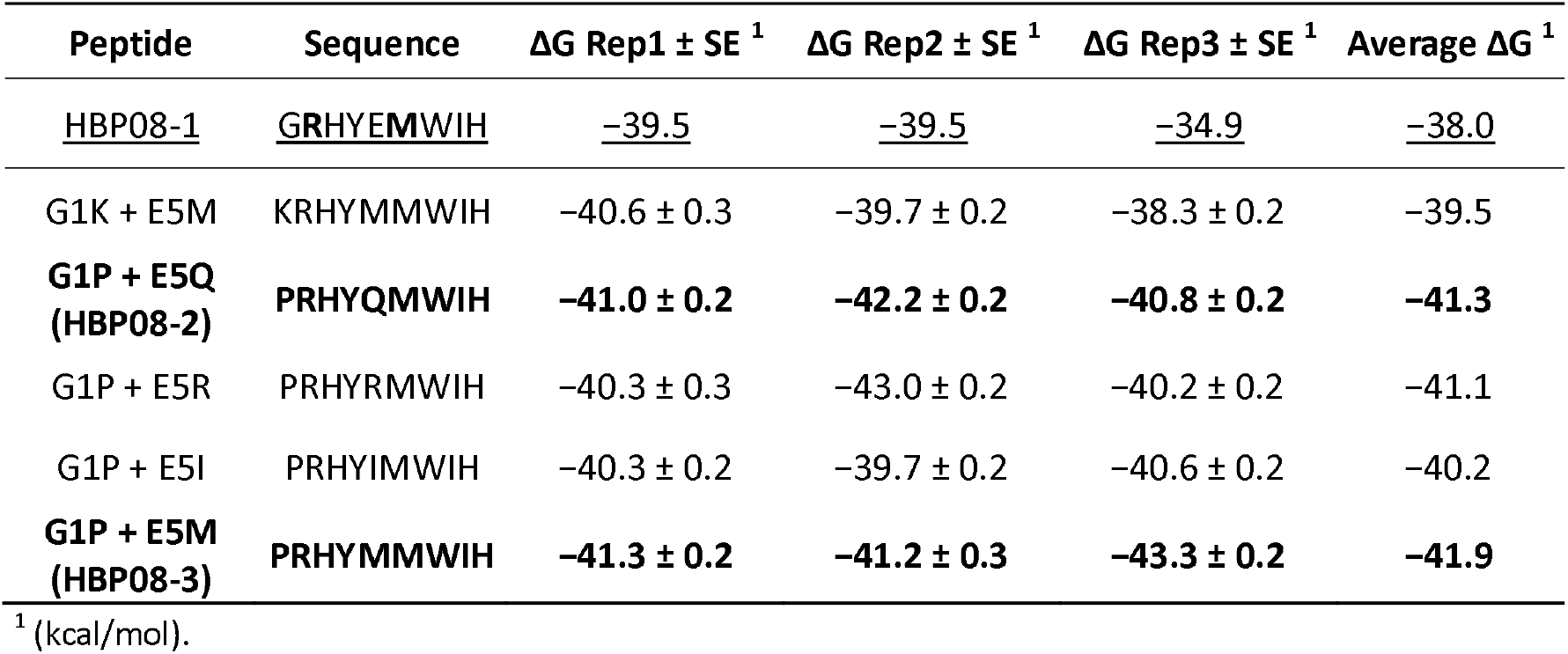
Binding free energy (ΔG) of the HBP08-1 mutant peptides in complex with HMGB1-BoxB, displaying ΔG values more than 2 kcal/mol lower to the parent peptide, subjected to two additional 500 ns MD simulations replicas.

The attained results highlighted that the mutant peptides G1P+E5Q (**HBP08-2**) and G1P+E5M (**HBP08-3**), which differ only by one residue in position 5, resulted in the most promising ones. Indeed, they showed an average ΔG value 3.3 and 3.9 kcal/mol lower compared to that of the parent HBP08 peptide, respectively (**Table 4**). Accordingly with these data, both peptides were purchased by Proteogenix (Schiltigheim, France) and then tested by MST experiments, using recombinant HMGB1, HMGB1-BoxA, and HMGB1-BoxB.

### Biophysical evaluation

The affinity of both **HBP08-2** and **HBP08-3** peptides for recombinant HMGB1-BoxB was assessed by MST as previously described (31), and displayed K_d_ values of 11.3 ± 2.3 nM and 15.3 ± 1.9 nM, respectively (**Figure 2AB**). Notably, these values are about 1000-fold lower compared to the one observed for HBP08 (17 µM) (31). Since both peptides displayed a similar K_d_ value, but **HBP08-3** had a lower solubility in PBS buffer, only **HBP08-2** was selected to perform MST experiments on recombinant HMGB1-BoxA and on the full length HMGB1, which revealed a K_d_ of 4.2 ± 0.4 µM and 28.1 ± 7.0 nM, respectively (**Figure 2CD**). Notably, the K_d_ value obtained assessing the affinity to full length HMGB1 further confirmed that the optimized **HBP08-2** peptide binds HMGB1 in the low nanomolar range, despite its reduced affinity to HMGB1-BoxA (**Figure 2C**).

**Figure 2.**
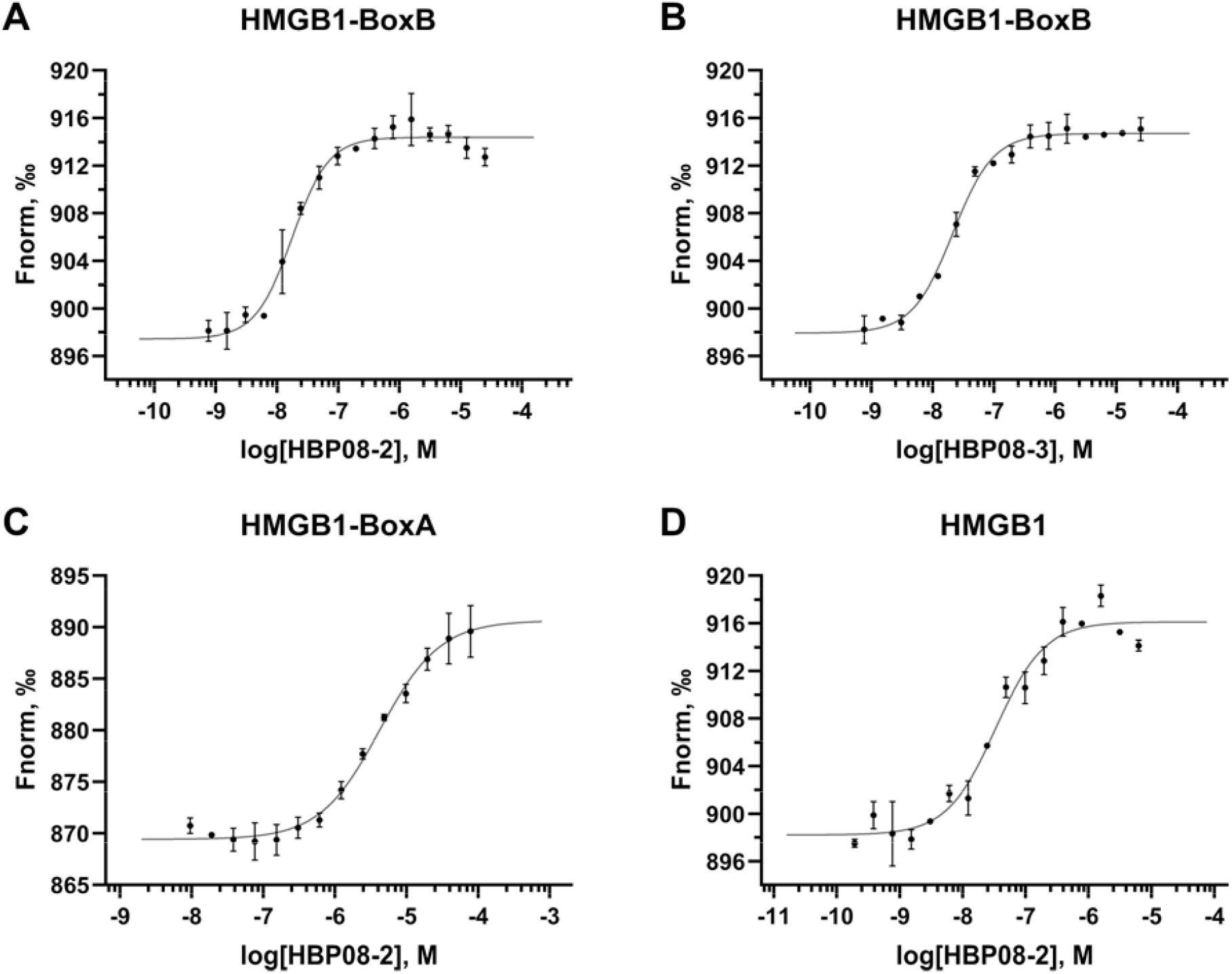
MST experiments. K_d_ curves of (A) HBP08-2 and (B) HBP08-3 on HMGB1-BoxB. (C) K_d_ curves of HBP08-2 on HMGB1-BoxA and (D) full sequence HMGB1.

### Biological evaluation

To evaluate the activity of the **HBP08-2** peptide on cell migration induced by the CXCL12/HMGB1 heterocomplex, we performed *in vitro* chemotaxis assays on primary human monocytes. **HBP08-2** abrogated the enhancement in cell migration induced by the CXCL12/HMGB1 heterocomplex, restoring migration to the levels induced by CXCL12 alone (**Figure 3A**). Notably, the newly designed **HBP08-2** peptide inhibited the heterocomplex activity at a concentration 10 times lower than that required for the parental HBP08 peptide, with an IC_50_ of 3.31 µM (**Figure 3B**). These findings were further confirmed using a murine cell line transfected with the human CXCR4, corroborating the improved potency of the **HBP08-2** peptide, which did not exhibit toxicity on either cell types (Supplementary Materials, **Figure S4-S5**). **HBP08-2** did not affect monocyte migration in response to CXCL12 alone, thus demonstrating the specificity of **HBP08-2** in inhibiting CXCL12/HMGB1-induced cell migration (**Figure 3C**).

**Figure 3.**
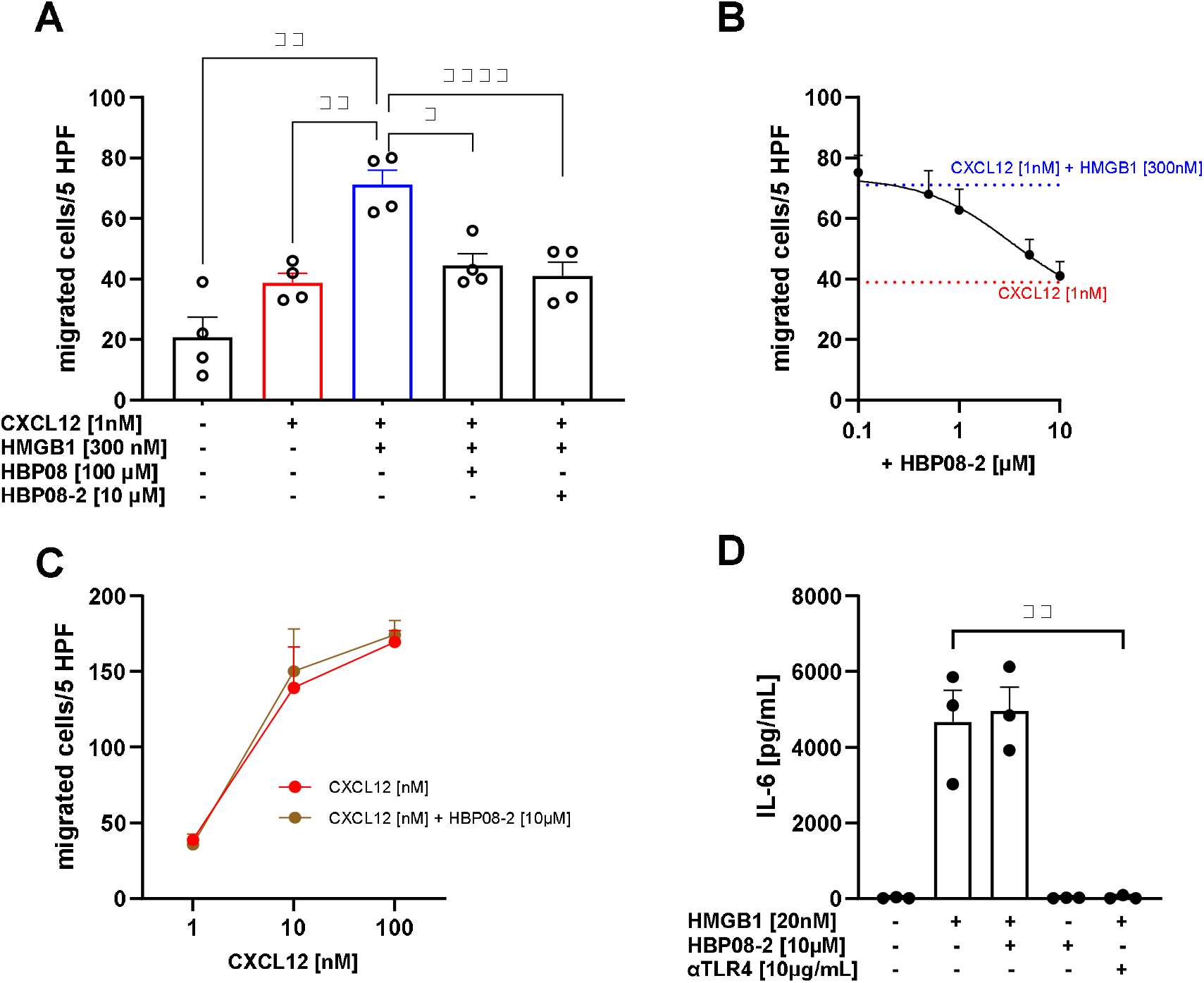
The HBP08-2 peptide selectively inhibits CXCL12/HMGB1 heterocomplex induced migration. (A) Inhibition of cell migration in response to the CXCL12/HMGB1 heterocomplex was assessed on human monocytes using the peptides HBP08-2 or HBP08 (as a control). *p < 0.05; **p < 0.01; ****p < 0.0001 by one-way ANOVA, followed by Dunnett’s multi-comparison test. (B) Inhibition of monocytes migration in response to the CXCL12/HMGB1 heterocomplex, assessed using scaling concentrations of the HBP08-2 peptide [10 μM, 5 μM, 1 μM, 0.5 μM and 0.1 μM], results in an IC_50_ of 3.31 µM. The blue line represents migration in response to the CXCL12/HMGB1 heterocomplex, while the red line represents migration in response to CXCL12 alone. (C) Monocytes migration in response to increasing concentration of CXCL12 in the presence or absence of the HBP08-2 peptide. (A-C) Migrated cells were counted in 5 high-power fields and data are shown as the mean + SEM of four independent experiments performed. (D) The concentration of IL-6 in the supernatant of monocytes treated with HMGB1 in the presence of the HBP08-2 peptide or of a neutralizing antibody against TLR4 (αTLR4) was measured by cytokine beads array. Data are shown as the mean + SEM of three independent experiments performed. **p < 0.01; by unpaired t test.

To investigate the activity of the **HBP08-2** peptide on HMGB1-mediated activation of TLR4, we measured cytokine release following stimulation of monocytes with recombinant HMGB1. The alarmin induced a significant release of IL-6 in the supernatant, which was effectively abrogated by adding a neutralizing antibody targeting TLR4 (**Figure 3D**). Importantly, **HBP08-2** neither hindered HMGB1-mediated cytokine release when added in combination with recombinant HMGB1, nor induced IL-6 release *per se*. These results underscore the selective inhibition by **HBP08-2** of the CXCL12/HMGB1 heterocomplex induced migration, allowing HMGB1 to maintain its TLR4-triggering capacity (**Figure 3D**).

### HBP08-2/HMGB1-BoxB model

In order to obtain a computational model of the **HBP08-2**/HMGB1-BoxB complex we performed a cluster analysis considering the three MD replicas (total simulation time = 1.5 µs), and the structure representative of the most populated cluster, which accounts for 96% of conformational ensembles explored (**Figure 4**). The side chains of R2 and Y4 of **HBP08-2** establish H-bond interactions with E131, while the carboxy-terminus group and the side chain of H9 establish H-bond and π-π interactions with R97 of HMGB1-BoxB, respectively, and I8 fits into the hydrophobic pocket surrounded by F102, A126, L129 and C106. Thus, **HBP08-2** exhibits a binding mode similar to that of **HBP08-1** (Supplementary Materials, **Figure S3B**). However, its slightly better predicted affinity (ΔG) and improved stability in the binding site during MD simulations may be attributed to the two mutations G1P and E5Q which, although their side chains are exposed to the solvent, allow the peptide to better interact with HMGB1-BoxB. In addition, the residue I8 of **HBP08-2** is positioned deeper in the hydrophobic pocket, establishing more hydrophobic interactions.

**Figure 4.**
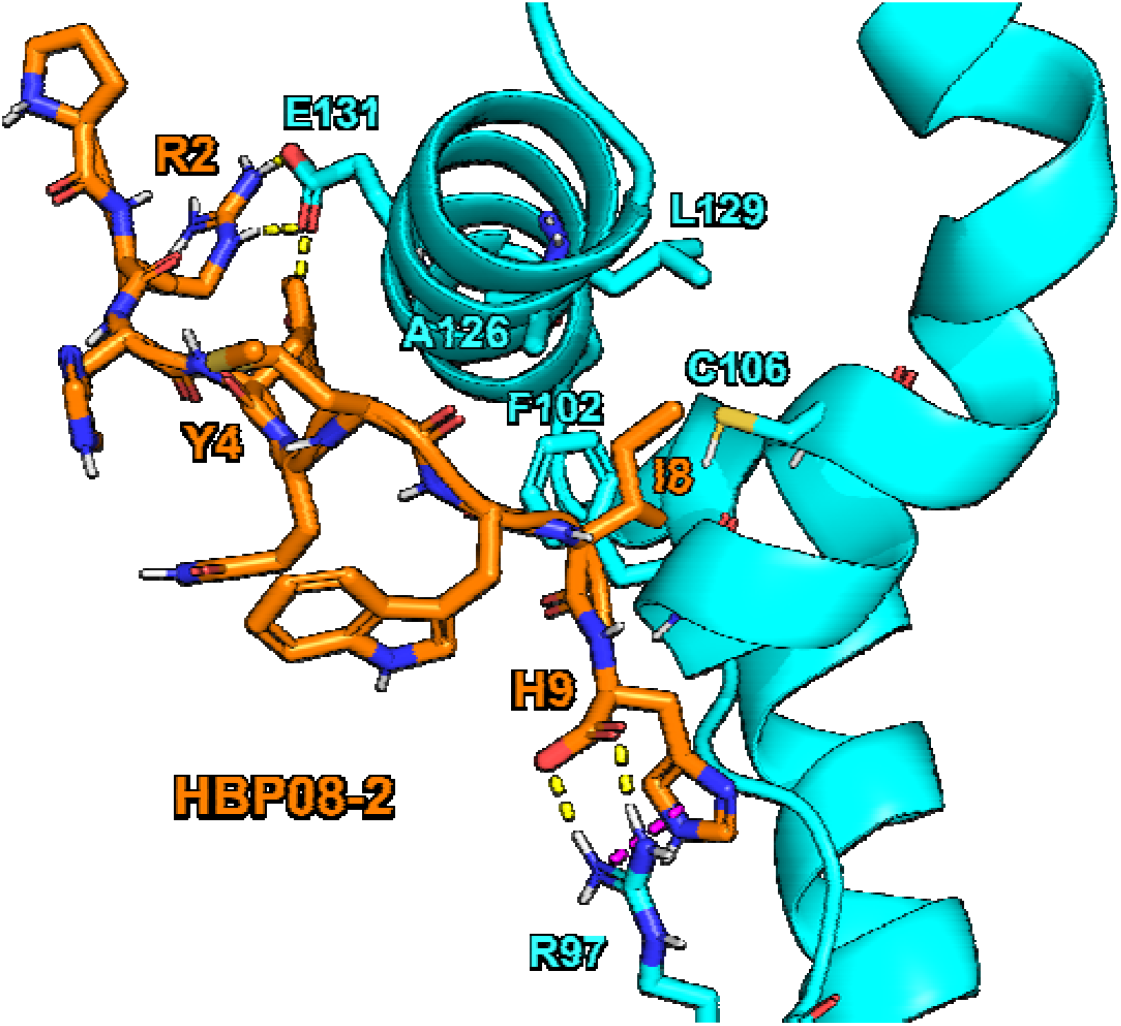
Representative structure of the most populated cluster of HBP08-2 (with sequence PRHYQMWIH, orange sticks) in complex with HMGB1-BoxB, considering 1.5 µs of MD simulations. H-bond and π-π interactions are represented as yellow and purple dashed lines, respectively.

## Materials & Methods

### HBP08/HMGB1-BoxB computational model

The HBP08/HMGB1-BoxB computational model used as starting point in this work was retrieved from our previous published article in which CSP NMR-guided docking calculations of HBP08 on HMGB1-BoxB were accomplished using Haddock (31,33). The initial protein was derived from the representative structure of the most populated cluster of fr-HMGB1 which was subjected to 30 × 1 µs independent MD simulations (total 30 µs) in a previous work from our group (17). Only the region from F90 to R163 was considered as the initial structure since it belongs to the BoxB domain (34). The HBP08/HMGB1-BoxB complex docking pose was then equilibrated through a long run MD simulation (500 ns) using Amber21 software.

### Molecular Dynamics (MD) simulations

MD simulations were performed in a system which was solvated in a TIP3P water box displaying a minimum distance of 10 Å from the protein surface, and counter-ions were added for charge neutrality. Amber force fields ff14SB (for protein atoms) (35), TIP3P model (for water molecules) (36), and parameters by Joung and Cheatham (for counter-ions) (37), were used to describe the system. To remove initial atom clashes, the system underwent a multi-step relaxation protocol involving energy minimization and gradual heating to 300 K over 300 ps, followed by an increase in pressure to 1 atm. Constant pressure and temperature were maintained using Monte Carlo barostat and Langevin thermostat, respectively (38,39). The SHAKE algorithm was applied to constrain hydrogen-involving bonds, and non-bonded van der Waals interactions were limited to a 9.0 Å cut-off range. Electrostatic interactions were treated with the particle mesh Ewald method (40). Simulations were run using the GPU-accelerated PMEMD code with a 2 fs time step, producing trajectories over 500 ns. RMSD calculations were accomplished using the VMD software (41), while the predicted binding affinity (ΔG) of the ligand was determined using the MMPBSA.py module available in Amber21 (42). In these calculations, the single trajectory approach was applied, and the entropy contributions to the binding free energy, coming from the normal-mode analysis, was neglected.

### Cluster analysis

The ligand conformations obtained from MD sampling were clustered using the Gromos method developed by Daura et al. (43), implemented in the GROMACS package (version 5.0.7) (44). After multiple clustering runs and accurate inspection of the results, we applied a proper RMSD threshold value to discriminate between the different ligand conformations while also limiting the amount of singleton clusters.

### Alanine scanning and affinity maturation protocol

Alanine scanning was performed by mutation of each residue of the native peptide sequence and each mutated peptide was subjected to a 500 ns-long MD simulation in complex with HMGB1-BoxB using the same procedure described above. Finally, MM-GBSA calculations were carried out to estimate the binding free energy (ΔG) of the mutated peptides to identify hot-spot residues. Non-hotspots residues were mutated into all possible combinations of natural amino acids by using the “Residue Scanning Calculation” tool, implemented in the BioLuminate module of Maestro (Schrödinger, LLC, New York, USA, version 2021-3) (45). The mutant peptides were ranked by ΔAffinity and ΔStability values, calculated by Prime MM-GBSA in implicit solvent. ΔAffinity refers to the change in binding affinity of the mutated peptide (ligand) to the HMGB1-BoxB (receptor), while ΔStability denotes the difference in free energy between the folded and unfolded states of the peptide due to the mutation. In both instances, negative values indicate that the mutant peptides exhibit better binding affinity or greater stability compared to the native protein. The best three mutant peptides by ΔAffinity, ΔStability and by considering both parameters (Mixed group) were subjected to MD simulations and ΔG estimation as described previously.

### Proteins and peptides

His-tagged HMGB1-BoxB, HMGB1-BoxA, and full length HMGB1 proteins were produced at the Protein Facility of the Institute for Research in Biomedicine as previously described (8,16), and stored in phosphate-buffered saline (PBS; product No. D8537, Sigma Aldrich, Saint Louis, USA). Both HBP08-2 and HBP08-3 peptides (without trifluoroacetic acid, TFA) were synthetized by Proteogenix (Schiltigheim, France), reconstituted with dimethyl sulfoxide (DMSO) for molecular biology (product No. D8418; Sigma-Aldrich, Saint Louis, USA), and stored at −20 °C. High-performance liquid chromatography (HPLC) and mass spectrometry (MS) confirmed that each peptide had a purity of about 99% (see Supporting Materials for the original documents).

### Microscale Thermophoresis (MST)

The binding affinity (K_d_) experiments between ligands (HBP08-2 and HBP08-3 peptides) and target proteins (HMGB1-BoxB, HMGB1-BoxA and full sequence HMGB1) were accomplished by using the Monolith NT.115^Pico^ instrument (NanoTemper Technologies GmbH, München, Germany). Target proteins were fluorescently-labeled using the specific His-Tag Labeling Kit RED-tris-NTA 2nd Generation of NanoTemper Technologies (Product No. MO-L018), according to manufacturer instructions. A fixed 10 nM concentration of labeled target protein was mixed with sixteen 1:1 serial dilutions of the ligand peptide (see Supplementary Materials, **Table S4** for the details about the concentration ranges used for the peptides). Both protein and peptide were dissolved in PBS-T buffer (0.05% Tween™ 20, NanoTemper Technologies), and incubated for at least 40 min at room temperature. MST analysis was conducted with premium-coated capillaries (product No. MO-K025; NanoTemper Technologies GmbH, München, Germany) applying an excitation and MST power of 20% and 40% (medium), respectively, at the fixed temperature of 25 °C. Before proceeding with the K_d_ determination, we ensured that the peptides were not auto-fluorescent. Finally, the data were processed by employing the dedicated MO.Affinity Analysis software v2.3 (NanoTemper Technologies GmbH, München, Germany) and the K_d_ values were determined based on concentration-dependent changes in normalized fluorescence (F_norm_), while the figures were generated using GraphPad Prism software v8.0.2 (GraphPad, Boston, USA). The dataset overview of the MST experiments accomplished is reported in the Supplementary Materials, **Table S4**.

### Cells

The murine 300.19 PreB cell line, stably transfected with the human CXCR4, was cultured under standard culture conditions (5% CO_2_, 95% O_2_, 37°C) in RPMI-1640, supplemented with 10% Fetal Bovine Serum, 1x non-essential amino acids, 1 mM Sodium pyruvate, 20 mM GlutaMAX, 50 μM β-Mercaptoethanol, 50 U/ml Penicillin and 50 μg/ml Streptomycin (GIBCO). Human monocytes were freshly isolated from buffy-coats obtained by spontaneous donation from healthy individuals (Central Laboratory of Swiss Red Cross, Basel, Switzerland, and Centro Trasfusionale Lugano, Switzerland, Canton Ticino Ethical Committee approval CE3428), and isolated by positive selection using CD14 microbeads (Miltenyi Biotec), as previously described (8).

### Chemotaxis Assay

*In vitro* cell migration of CXCR4+ murine 300.19 PreB cells and of freshly isolated human monocytes was performed using Boyden chambers equipped with 5 μm pore membranes, as previously described (46). Briefly, cells were diluted at 10^6^ cell/mL in RPMI 1640, supplemented with 20 mM Hepes, pH 7.4, and 1% pasteurized plasma protein solution, and allowed to migrate for 90 min in response to different stimuli under standard conditions (5% CO_2_, 95% O_2_, 37 °C). CXCL12 was used at 1 nM, 10 nM or 100 nM. The CXCL12/HMGB1 heterocomplex was formed by combining 1 nM CXCL12 with 300 nM HMGB1. To assess the inhibitory effect of the **HBP08-2** peptide, different concentrations [0.1 μM, 0.5 μM, 1 μM, 5 μM, and 10 μM] were incubated with the CXCL12/HMGB1 heterocomplex. The HBP08 peptide at 100 μM, was used as positive control (31).

### Assessment of Peptide Toxicity

**HBP08-2** toxicity was assessed on the murine CXCR4+ 300.19 PreB cell line and on freshly isolated human monocytes, as described previously (31). Briefly, cells were cultured for 2 or 4 hours in the presence of the peptide HBP08 or **HBP08-2** at 1 μM or 10 μM. After incubation, cells were stained using AnnexinV-FITC and propidium iodide following the manufacturer’s instructions, and cell viability was analyzed by flow cytometry in comparison to the untreated control.

### Cytokine Quantification

Freshly isolated human monocytes were incubated for 8 hours under standard conditions (5% CO_2_, 95% O_2_, 37 °C) at 10^6^ cell/mL in RPMI-1640 supplemented with 0.05% pasteurized human albumin in the presence or absence of 20 nM HMGB1. To inhibit TLR4 activation, a neutralizing antibody was used at 10 μg/mL (AF1478, R&D System). The **HBP08-2** peptide at 10 μM was tested for its ability to inhibit HMGB1-mediated release of cytokines. Quantification of IL-6 in the supernatants was determined by using the cytometric bead array (CBA)-Human Inflammatory Cytokines Kit (551811, BD Biosciences, San Jose, CA, USA). Acquisition was performed with a FACS Canto II (BD Biosciences, San Jose, CA), and the cytokine concentration was calculated from the mean fluorescence intensity according to a standard curve for IL-6.

## CONCLUSION

The application of a computational approach led us to optimize the HBP08 sequence for obtaining a mutated peptide (namely **HBP08-2**) with improved binding affinity to HMGB1-BoxB and full length HMGB1. Therefore, **HBP08-2** peptide represents the HMGB1 binder displaying the highest affinity reported in literature so far.

Functional *in vitro* assays, aimed at characterizing the biological activity of **HBP08-2**, showed its selectivity in inhibiting CXCL12/HMGB1 heterocomplex-induced cell migration, with an IC_50_ which is about ten times lower compared to the one observed for its parent peptide HBP08 (31). Importantly, **HBP08-2** exhibited no interference with either CXCL12-induced migration or HMGB1-mediated TLR4 activation. These additional findings strengthen the unique pharmacological profile of **HBP08-2**.

**HBP08-2** can serve as a valuable tool for cell biologists to investigate the inflammatory pathways initiated by the CXCL12/HMGB1 heterocomplex, *in vitro* and *in vivo*. We believe that the structural information provided in this work on the **HBP08-2**/HMGB1-BoxB complex could be useful for the development of new peptidomimetics, with improved pharmacokinetic profiles, which could represent a new class of anti-inflammatory drugs. In particular, it could be useful for those patients, poorly responding to current therapies, affected by chronic inflammatory conditions, such as rheumatoid arthritis, in which the pro-inflammatory role of the CXCL12/HMGB1 heterocomplex is crucial (21).

## Author Contributions

E.M.A.F. and E.P. contributed equally to this work. E.M.A.F. designed, performed, and analyzed the computer simulations, performed MST experiments and wrote the manuscript. E.P. performed the *in vitro* experiments and wrote the manuscript. A.C. discussed the results of both experimental and computational studies, wrote the manuscript. G.R. supervised the work and wrote the manuscript. V.C. and M.U. designed the *in vitro* experiments, supervised the work and wrote the manuscript. J.S and G.G. designed the computational pipeline, supervised the work and wrote the manuscript. All the authors discussed, reviewed, and approved the manuscript.

## Notes

The authors declare the following competing financial interest(s): A.C., M.U. and J.S. are holding a patent entitled “PEPTIDE INHIBITORS TARGETING THE CXCL12/HMGB1 INTERACTION AND USES THEREOF”, with priority date 21 March 2019, publication number WO2020188110.

## Acknowledgement

This study was supported by the Swiss National Science Foundation (3100A0-143718/1 to M.U.). Special thanks to Gabriela Danelon for her excellent technical support to our work. G.G. would like to thank INDACO for providing high-performance computing resources and support.

