## Supplementary Material for "A Potent Nonapeptide Inhibitor for the CXCL12/HMGB1 heterocomplex: A Computational and Experimental Approach"

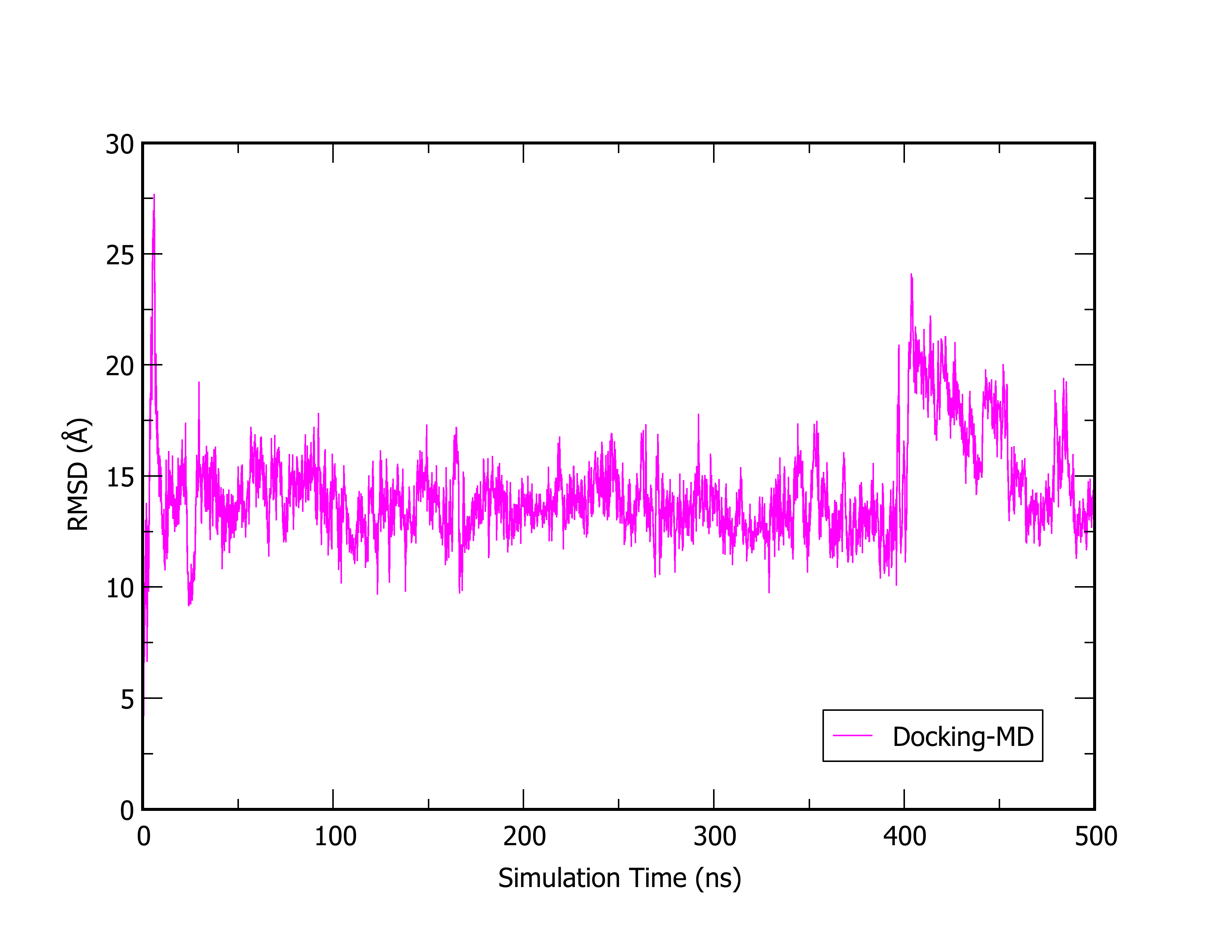

**Figure S1.** RMSD analysis of the HBP08/HMGB1-BoxB complex model subjected to 500 ns-long MD simulation. The complex structure was retrieved in our previous work published (DOI: 10.1021/acs.jmedchem.1c00852), that was obtained by computational docking guided by the protein residues involved in the peptide binding as reported by NMR CSP experiments performed.

**Table S1**. Binding free energy (ΔG) values of each independent replica of the HBP08/HMGB1-BoxB complexes.

| **HBP08** | **ΔG ± SE (kcal/mol)** |
| --- | --- |
| MD replica1 | -31.9 ± 0.4 |
| MD replica2 | -31.2 ± 0.3 |
| MD replica3 | -36.4 ± 0.5 |

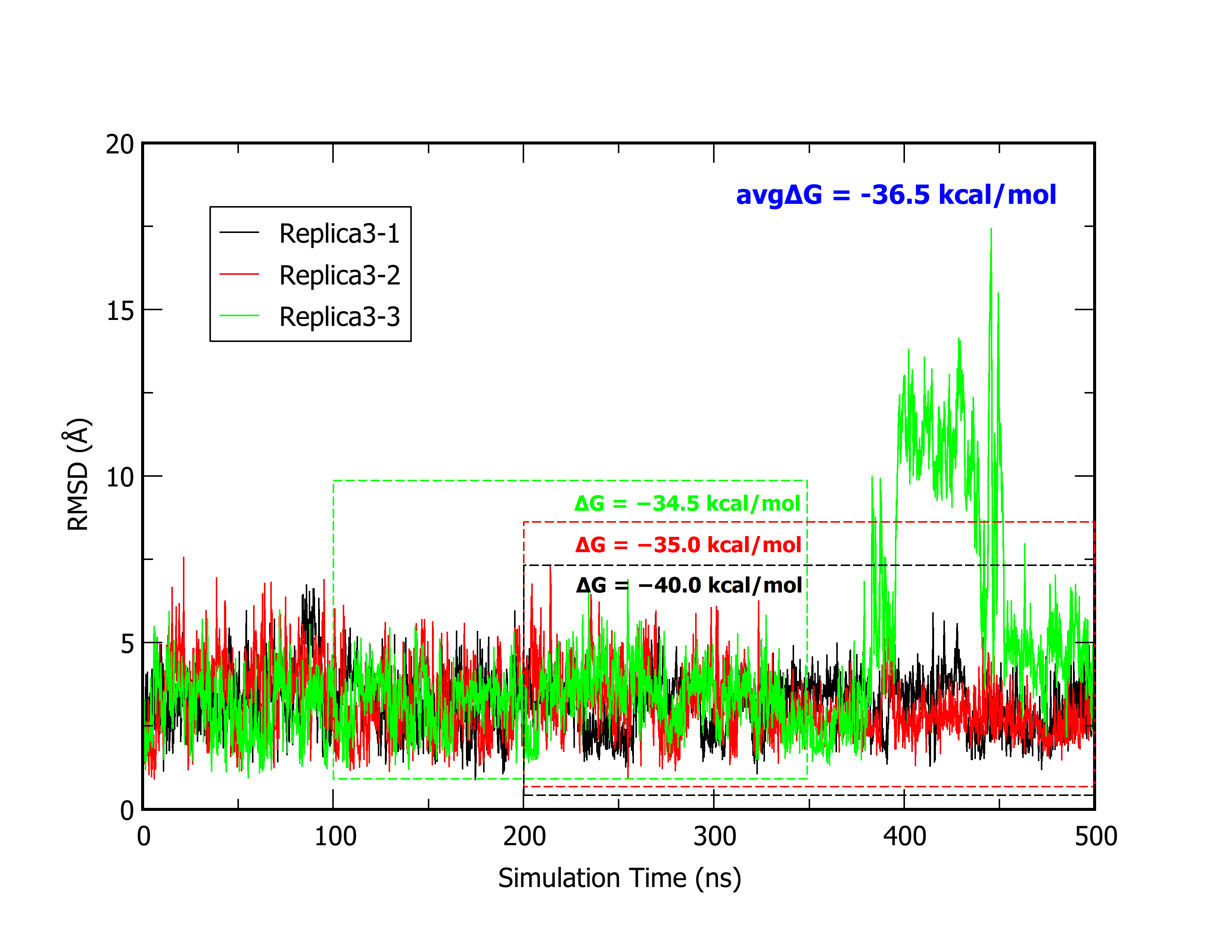

**Figure S2.** RMSD analysis of replica3-1 (black), replica3-2 (red) and replica3-3 (green) of HBP08 in complex with HMGB1-BoxB protein. In broken lines are highlighted the snapshots considered in the MM-GBSA calculations. The ΔG values obtained for each replica and the average value are reported.

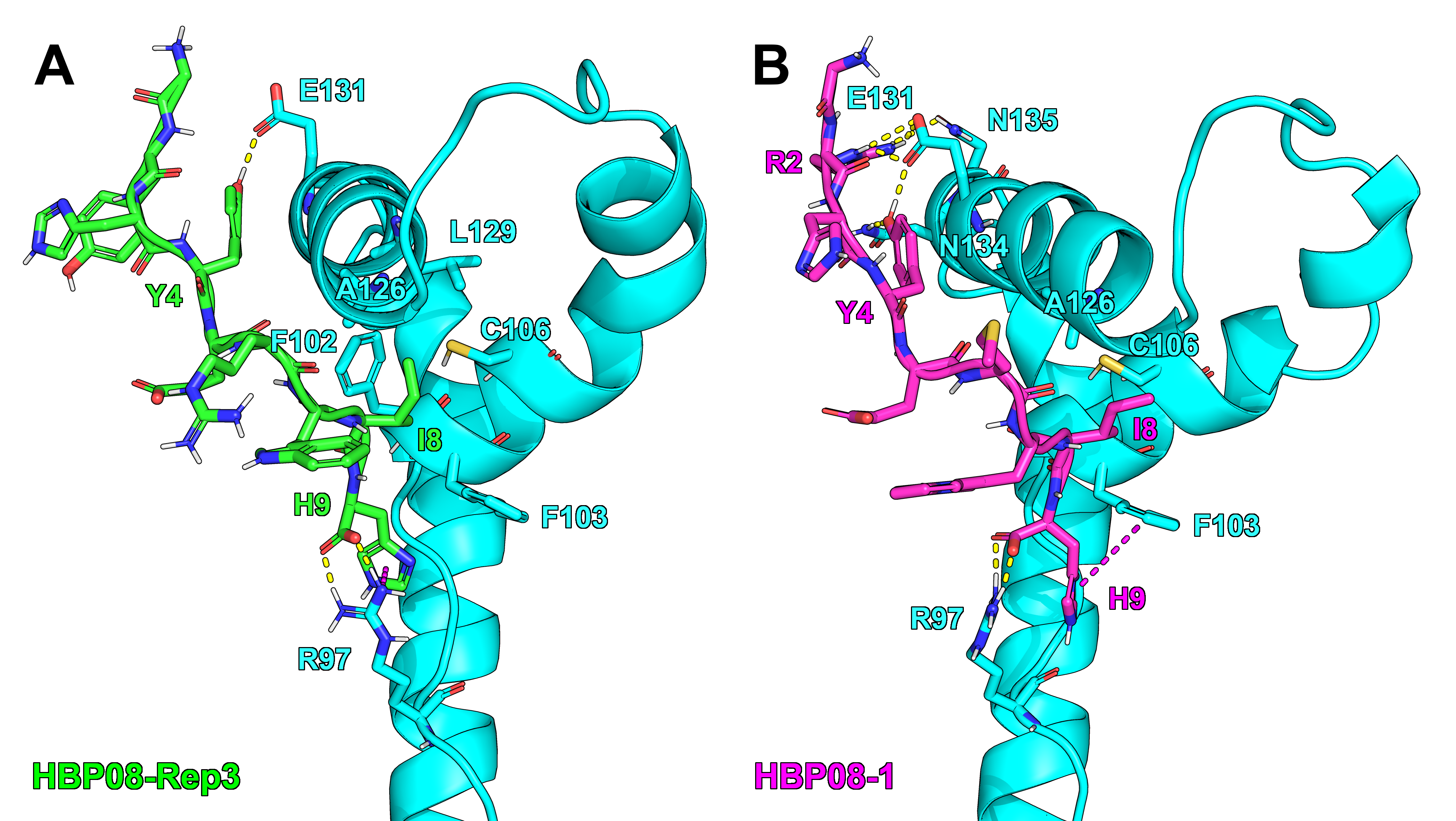

**Figure S3.** Representative structure of the most populated cluster of (A) HBP08-Replica3 (with sequence GYHYERWIH, green sticks) and (B) HBP08-1 peptide (with sequence GRHYEMWIH, magenta sticks) in complex with HMGB1-BoxB, considering 1.5 µs of MD simulations. These complexes were used as starting complex structure for the following affinity maturation steps. H-bond and π-π interactions are represented as yellow and purple dashed lines, respectively.

**Table S2**. Binding free energy (ΔG) of the mutated peptide HBP08-1 compared to the parent peptide HBP08.

| **Peptide** | **Sequence** | **ΔG Rep1 ^1^** | **ΔG Rep2 ^1^** | **ΔG Rep3 ^1^** | **Average ΔG ^1^** |
| --- | --- | --- | --- | --- | --- |
| HBP08 | GYHYERWIH | -40.0 | -35.0 | -34.5 | -36.5 |
| HBP08-1 | G**R**HYE**M**WIH | -39.5 | -39.5 | -34.9 | -38.0 |

^1^ (kcal/mol).

**Table S3**. Binding free energy (ΔG) of the HBP08-1 mutated peptides in complex with HMGB1-BoxB, derived from the affinity maturation protocol in which H3 and I8 were simultaneously mutated, subjected to 500 ns MD simulations.

| **Group** | **Mutation** | **Sequence** | **ΔAffinity ^1^** | **ΔStability ^1^** | **ΔG ± SE ^1^** |
| --- | --- | --- | --- | --- | --- |
| **HBP08-1** | / | G**R**HYE**M**WIH | / | / | −38.0 |
|  | H3E + I8M | GREYEMWMH | −12.19 | −2.26 | −37.5 ± 0.3 |
| **Affinity** | H3Y + I8M | GRYYEMWMH | −11.64 | 0.00 | −35.4 ± 0.3 |
|  | H3N + I8M | GRNYEMWMH | −9.92 | +4.08 | −36.5 ± 0.2 |
|  | H3R + I8L | GRRYEMWLH | +3.09 | −13.56 | −35.9 ± 0.3 |
| **Stability** | H3R + I8G | GRRYEMWGH | +19.01 | −13.35 | *unbound* |
|  | H3R + I8M | GRRYEMWMH | −7.32 | −13.14 | −37.5 ± 0.2 |
| **Mixed** | H3Q + I8M | GRQYEMWMH | −8.88 | −4.54 | −34.2 ± 0.3 |
|  | H3L + I8M | GRLYEMWMH | −8.39 | −5.44 | −32.8 ± 0.3 |

^1^ (kcal/mol).

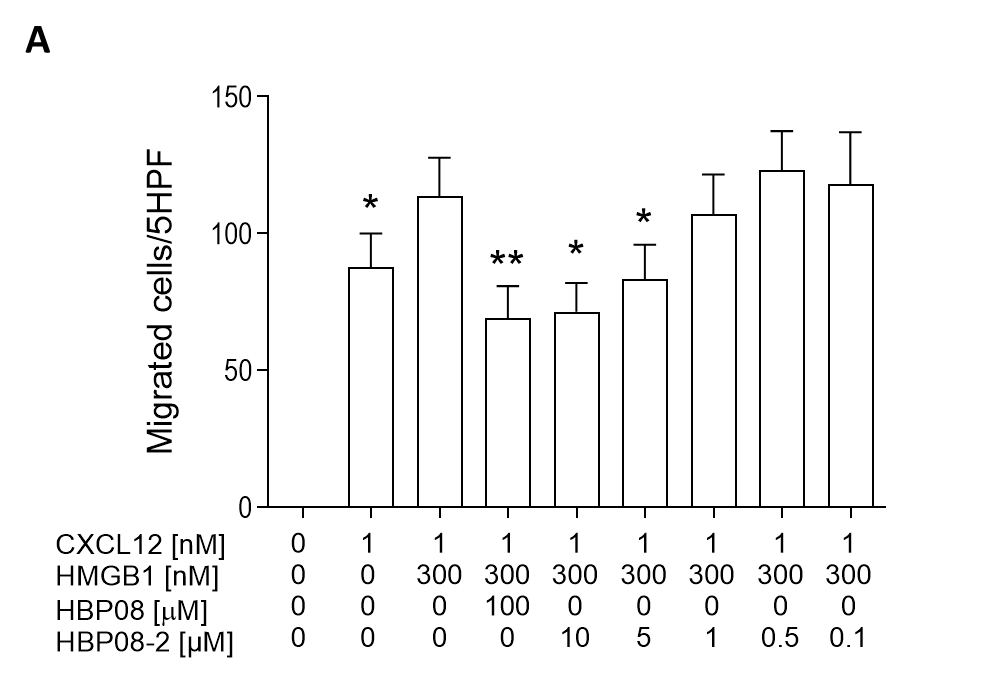

**Figure S4**. *In vitro* inhibition of CXCR4-transfected 300-19 Pre-B cells in response to the CXCL12/HMGB1 heterocomplex using the HBP08-2 peptide. (A) Inhibition of cell migration in response to the CXCL12/HMGB1 heterocomplex was assessed on 300-19 Pre-B cells CXCR4-transfected using the identified peptide HBP08-2 or HBP08 (as a control). Data are shown as the mean + SEM of four independent experiments performed. *p < 0.05; **p < 0.01; by one-way ANOVA, followed by Dunnett’s multicomparison test, comparing each condition to the migration observed in response to the CXCL12/HMGB1 heterocomplex.

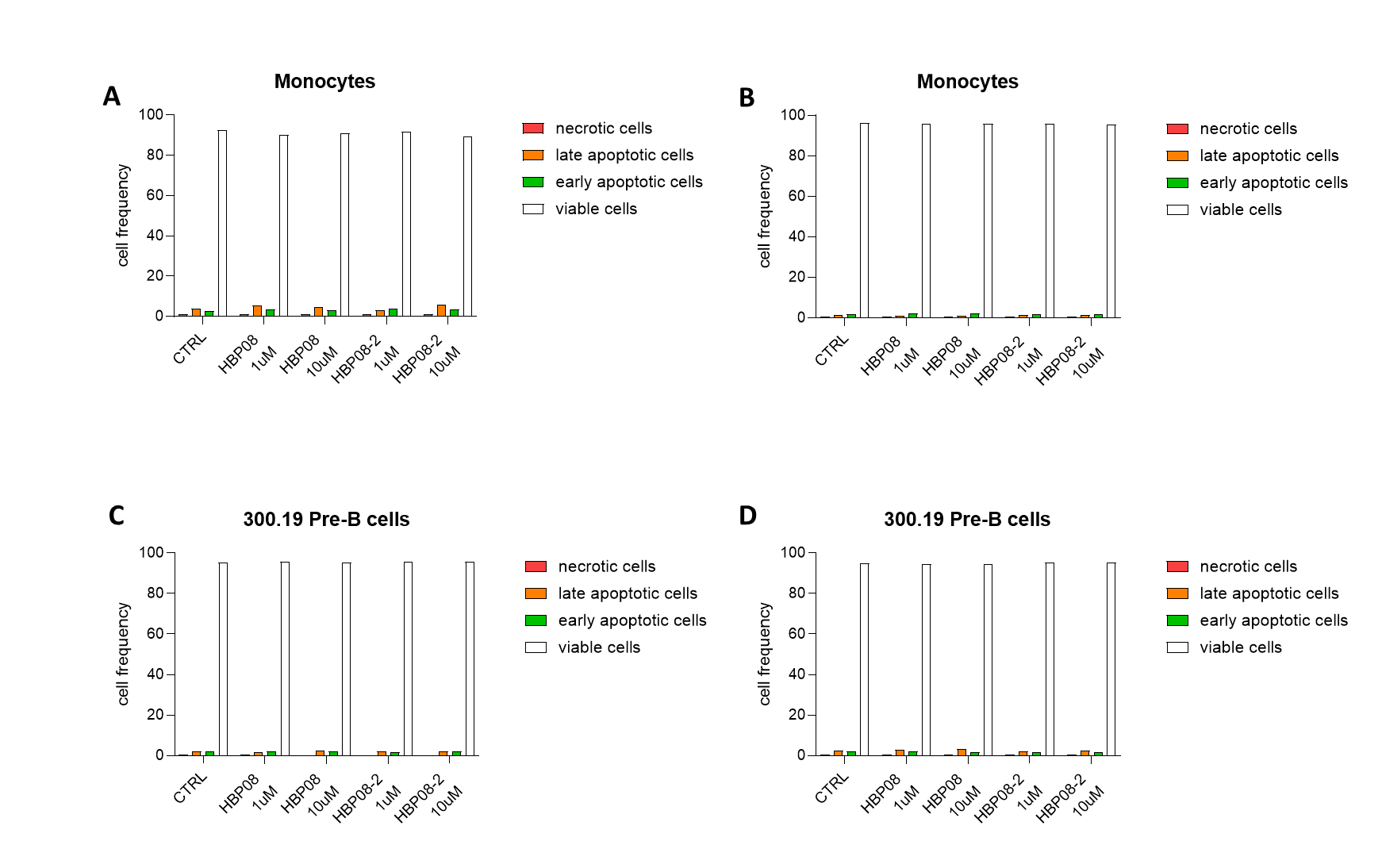

**Figure S5.** *In vitro* determination of cell viability by Annexin V/PI staining and flow cytometric analysis. Monocytes (A-B) and 300.19 Pre-B cells transfected with CXCR4 (C-D) were treated with 1 or 10 μM of HBP08 or HBP08-2 for 2 hours (A-C) or 4 hours (B-D).

**Table S4.** Dataset overview of the Microscale Thermophoresis (MST) experiments accomplished using a fixed 10 nM concentration of target proteins (HMGB1-BoxB, HMGB1-BoxA and full sequence HMGB1) and different concentrations of HBP08-2 and HBP08-3 peptides. At least two independent experiments were performed to compute the K_d_ value.

| **Peptide** | **Target Protein** | **MST Power** | **Exc. Power** | **Temp.** | **[Ligand] Range** | **Time** | **RA** | **SNR** | **K_d_ (nM)** |
| --- | --- | --- | --- | --- | --- | --- | --- | --- | --- |
| HBP08-2 | HMGB1-BoxB | 40% | 20% | 25 °C | 25 µM – 0.763 nM | 5 s | 17.8 | 20.1 | 11.3 ± 2.3 |
| HBP08-3 | HMGB1-BoxB | 40% | 20% | 25 °C | 25 µM – 0.763 nM | 5 s | 17.4 | 31.9 | 15.3 ± 1.9 |
| HBP08-2 | HMGB1-BoxA | 40% | 20% | 25 °C | 78.10 µM – 9.54 nM | 5 s | 21.4 | 36.6 | 4243 ± 447 |
| HBP08-2 | HMGB1 | 40% | 20% | 25 °C | 6.25 µM – 0.19 nM | 5 s | 18.1 | 14.2 | 28.1 ± 7.0 |

RA = Response Amplitude; SNR = Signal-to-Noise Ratio.
